# Discovery of μ, δ-Opioid receptor dual biased agonists that overcome the limitation of prior biased agonists

**DOI:** 10.1101/2021.01.21.425817

**Authors:** Jin Hee Lee, Suh-Youn Shon, Woojin Jeon, Sung-Jun Hong, Junsu Ban, Do Sup Lee

## Abstract

Morphine is widely used to manage pain in patients, although the risk of side effects is significant. The use of biased agonists to the G protein of μ-opioid receptors has been suggested as a potential solution, although Oliceridine and PZM21 have previously failed to demonstrate benefits in clinical studies. An amplification-induced confusion in the process of comparing G protein and beta-arrestin pathways may account for previous biased agonist mis-identification. Here, we have devised a strategy to discover biased agonists with intrinsic efficacy. We computationally simulated 430,000 molecular dockings to the μ-opioid receptor to construct a compound library. Hits were then verified by experiment. Using the verified compounds, we performed simulations to build a second library with a common scaffold, and selected compounds which show biased features to μ and δ-opioid receptors through a cell-based assay. Three compounds (ID110460001, ID110460002, and ID110460003) with a dual biased agonistic effect for μ and δ-opioid receptors were identified. These candidates are full agonists for the μ-opioid receptor, and they show specific binding modes. Based on our findings, we expect our novel compound to act as a biased agonist than conventional drugs such as Oliceridine.

## Introduction

Morphine, the archetypal analgesic opioid, is broadly used to manage pain due to external injury or underlying disease. Morphine and opioid-derived drugs target the opioid receptor family, including μ-(OPRM), δ-(OPRD), and κ-(OPRK) receptors. Opioid receptors are G protein-coupled receptors (GPCR), and are activated through G protein subunits and β-arrestin. Although it demonstrates superior efficacy to other analgesics, morphine can also induce several adverse effects.To overcome the side effects of morphine, agonists that target a subgroup of opioid receptors^1,2^, or only peripheral opioid receptors ^3^, have been developed. In another approach, agonists showing functional selectivity have been exploited^4–6^.

Although opioid receptors are activated by the G protein and β-arrestin2 pathways, these pathways do not contribute equally to the pain relief effect. As demonstrated by *in vivo* experiments using β-arrestin2 knockout rodents, the analgesic effect of morphine is predominantly determined by G protein, while the side effects were determined by β-arrestin2^7^. Thus, research has focused on G protein functional selective agonists ^8^. Oliceridine, an apparent “biased agonist” targeting OPRM, has already been evaluated in preclinical studies and clinical trials ^4,9–12^. However, Oliceridine was only approved by the US FDA as a full agonist of OPRM, essentially because it did not show significantly reduced respiratory depression in clinical trials. PZM21, another biased agonist against OPRM, was discovered using a structure-based docking simulation^5^. In cell-based assays, PZM21 demonstrated better potency and efficacy than Oliceridine. Subsequent animal experiments confirmed the analgesic effects of PZM21, and provided evidence for reduced side effects, including respiratory depression. However, subsequent studies have reported that PZM21 is a partial agonist for OPRM^13^, and clinical benefits are limited ^14^.

Two different approaches have been used to calculate biased agonist features such as functional selectivity: “Equiactive comparison ^15^”, used in the discovery of Oliceridine; and the “Black-Leff operational model with transduction coefficient ^16^”, used in PZM21 profiling. In the equiactive comparison method, the respective concentrations of agonists in the two pathways are assumed from each concentration-response curve. Using this approach, a bias factor (including the signaling state of relative activity) is obtained. In the Black-Leff operational model, because agonist binding initiates downstream signaling pathways, the efficacy of the receptor pathway can be defined by receptor occupancy populations. The operational model is described by the τ parameter index, which defines the intrinsic coupling efficiency of the agonist, and the logK_A_ parameter, which denotes the functional equilibrium dissociation constant of the agonist ^17^. Using these two parameters, the logarithm of the “transduction coefficient” (τ/K_A_) parameter can be calculated. The difference in transduction coefficient parameter of each pathway is calculated to determine the relative activities of the agonist in the different signaling pathways. For this procedure, test compound data is compared to full agonist data, a reference criteria of maximal receptor signaling efficacy ^18^. By using a reference agonist, data variance from environmental factors is reduced. Hence, the transduction coefficient method was the standard for bias calculation. However, PZM21 did not show promising preclinical study despite using the transduction coefficient methods^13^. Thus, there is a disconnect between cell-based assay results and *in vivo* systems.

After receptor-ligand binding, physiological signaling pathways are initiated, and this signaling becomes more amplified the further downstream^19^. The amplification is mediated by intracellular molecules such as Ca^2+^ and cyclic adenosine monophosphate (cAMP)^20^. Comparative activation of most existing biased agonists for opioid receptors was conducted using intracellular cAMP measurements for G protein pathway activation and β-arrestin2 recruitment to the intracellular receptor region for β-arrestin2 pathway activation. However, results from this comparison may overestimate the G protein efficacy due to unbalanced signaling amplification. Furthermore, the cell lines used in the *in vitro* assay systems are genetically engineered to overexpress target receptors. Therefore, these cell lines have a higher population of receptors compared with physiological systems. Thus, *in vitro* experiments have a high receptor reserve, and most agonists tend to maximal efficacy (like full agonists) under these conditions^21^. Under conditions where signaling amplification and receptor reserve exist simultaneously, partial agonists may behave as biased agonists that reach maximal effects in the G protein pathway^22^. By utilizing a lower receptor population and comparing the G protein pathway and β-arrestin pathway at the same signaling distance, Alexander Gillis et al^23^ revealed that Oliceridine, PZM21, and SR-17018 are partial agonists. To discover true biased agonists, it is necessary to confirm that ligands in different signaling cascades within the *in vitro* system are unaffected by the abovementioned limitations.

To discover a new scaffold for opioid receptor biased agonists, we applied a virtual screening (VS) methodology. This is a widely used technique for accelerating the early drug discovery process and identifying novel scaffolds. In many studies, VS is significantly faster and equally or more successful than high-throughput screening (HTS). Moreover, VS costs around 10-fold less than HTS and takes approximately half the time^24^.

Here, we carried out a sequential ligand-based and structure-based virtual screening approach (Figure 1). The Korea Chemical Bank (KCB) was preprocessed using Pipeline Pilot ^24^. The virtual hits were then screened to discover OPRM, OPRD dual biased agonists for the G protein pathway. To minimize uncertainty due to the limitations of signal amplification and receptor reserve, we used cell lines in which receptor population can be tuned by irreversible antagonists. Thus, we obtained compounds with a bias effect emanating from high efficacy. The binding sites of our compounds were subsequently evaluated to improve the efficacy profiles.

**Figure 1.**
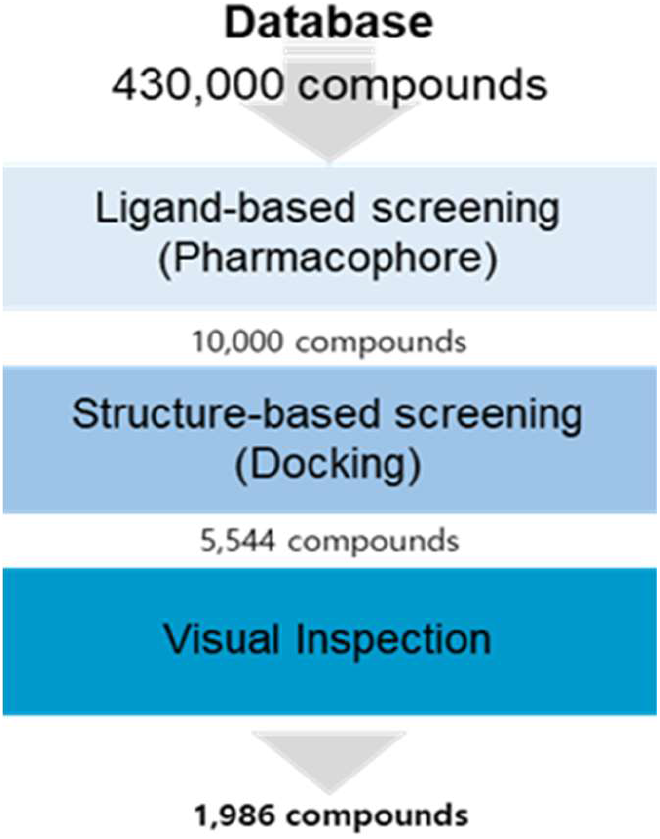
Virtual screening workflow.

## Results and Discussion

An efficient strategy for screening chemical candidates for OPRM, OPRD dual biased agonists was devised. We adopted a virtual screening method using the KCB database and validated the results using the cell-based assay. Using chemistry aided drug discovery (CADD) and the cell-based assay we were able to define a common chemical structure. This was then applied to the docking simulation. We identified three candidates with different mechanisms of action to Oliceridine and PZM21. The bases of these differences were then explained through simulations of molecular models.

### Virtual screening

We applied pharmacophore-based virtual screening using the reported OPRM biased agonists. The lowest energy conformations of Oliceridine, PZM21, and SR-1708 were geometrically refined using Discovery studio. These structures were used to build a three-dimensional common feature model within Phase. The resulting four-feature pharmacophore comprised one hydrophobic group, two aromatic rings, and one positive ionic feature. Figure 2 shows the three reported biased agonists superimposed on these features and especially, their protonated amine group was mapped the positive ionic feature (Figure 2). The pharmacophore model was then applied as a filter to virtually screen the KCB library. The top 10,000 candidates (by score) were selected for the docking study. To narrow the focus, molecular docking was performed upon active state OPRM using Glide SP. The reported biased agonist binding modes demonstrated docking scores less than −5.9. This value was set as the cutoff to be used for filtering in our structure-based virtual screening (SBVS). In total, 5,544 unique compounds remained following the pharmacophore-based and docking-based filtering stages. To exclude false-positive compounds, we performed clustering with ECFP4 and an additional visual inspection within the binding site of OPRM. Finally, we identified 1,986 potential compounds for biological testing.

**Figure. 2.**
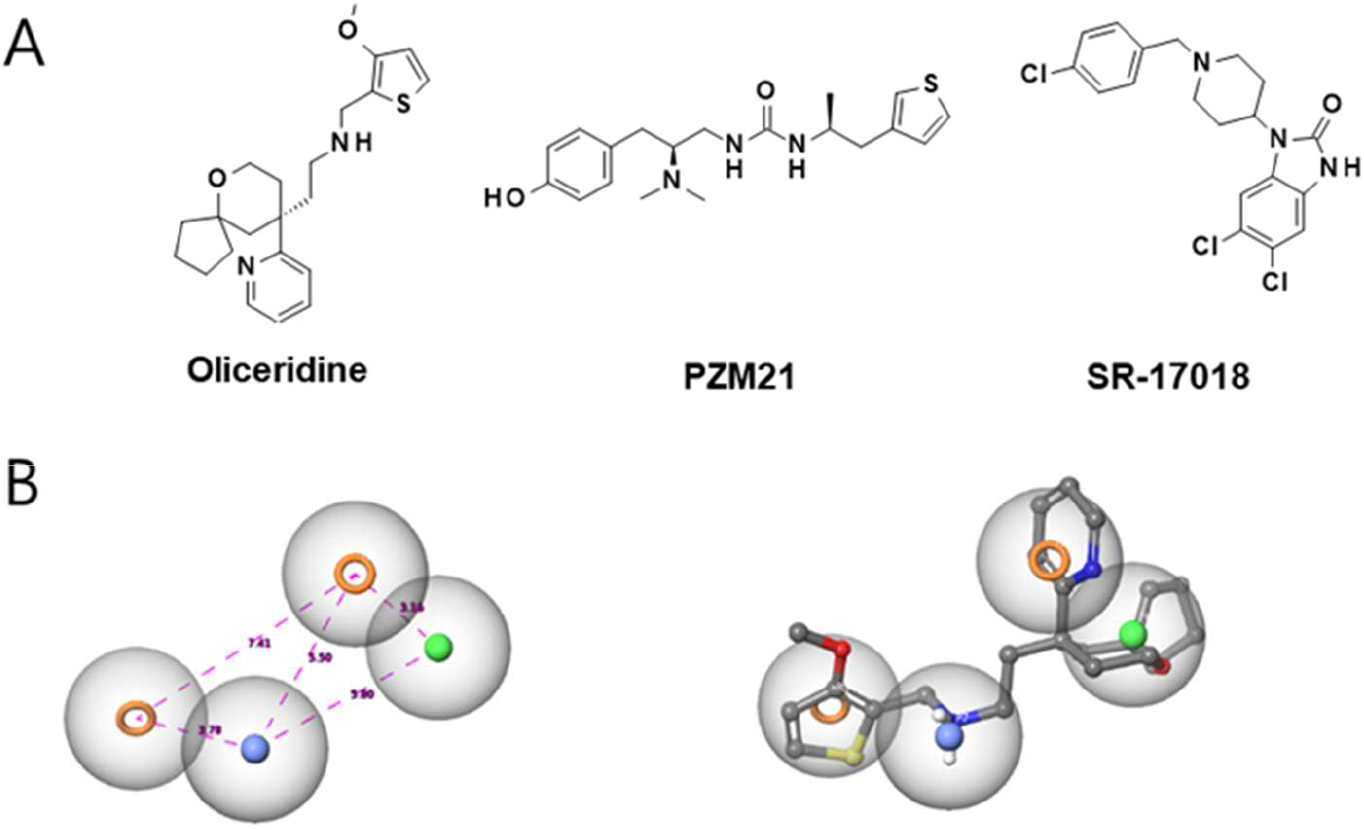
Pharmacophore model. (A) Chemical structures of the three biased agonists. (B) The four-features pharmacophore model (with Oliceridine): Green sphere, hydrophobic region(H); orange rings, aromatic ring feature (RA); dark-blue, positive ionic feature (P).

### Screening of compounds at defined concentrations

The flow chart for the discovery of OPRM, OPRD dual biased agonists is described in Figure 3A. The previously identified set of small molecules was screened through a cAMP assay to evaluate their activation properties. The first screening was accomplished at a single concentration (10±1 μM). Because virtual screening was focused on OPRM, the primary target in this screening was OPRM. As described in Figure 3B, 55 compounds showed at least 50% activation when compared with the E_max_ of DAMGO. These compounds were screened again in triplicate at the same concentration, and the results were reproducible for 32 compounds (Figure 3C).

**Figure 3.**
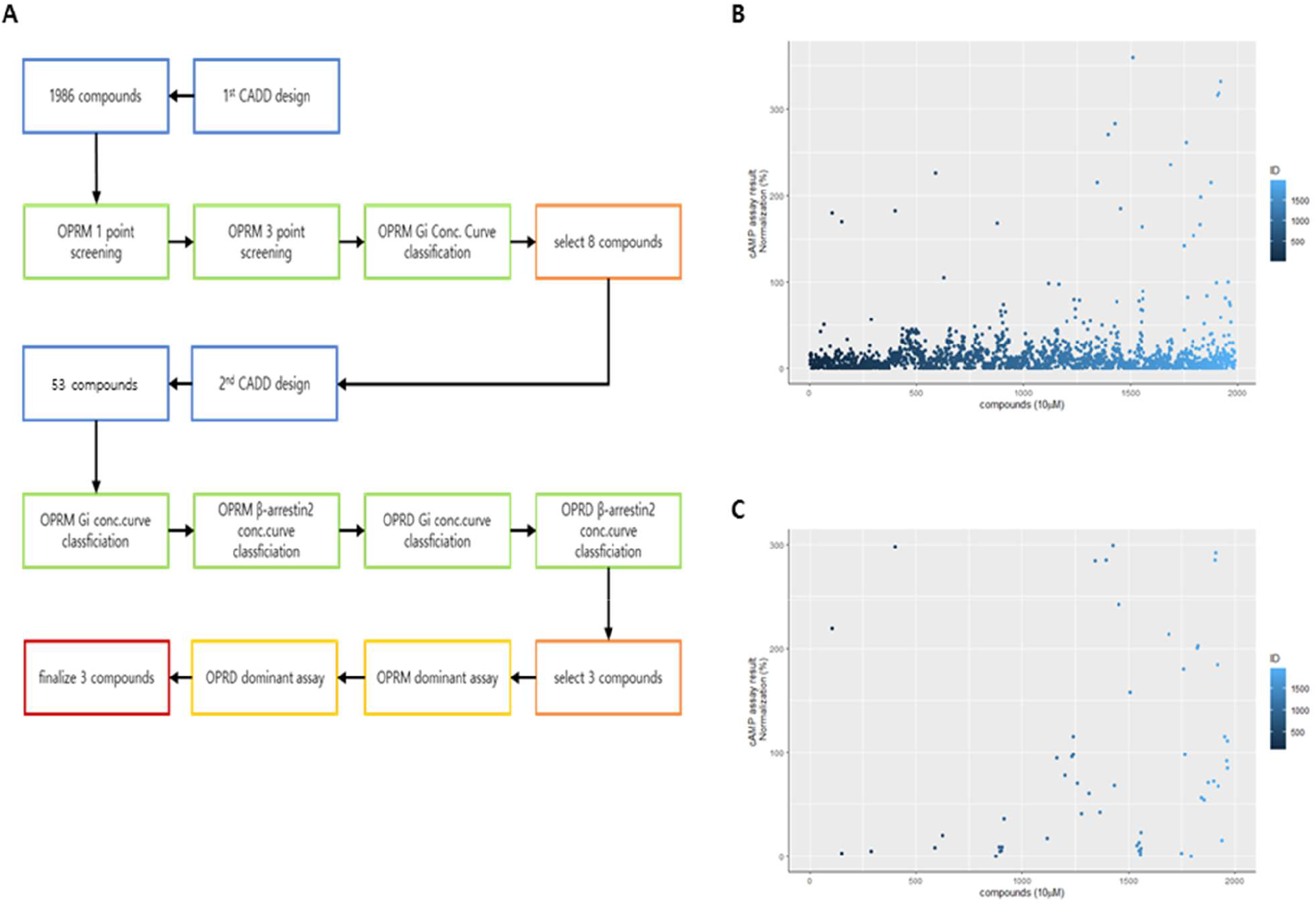
Screening results for 1986 compounds and assay flowchart. (A) Flowchart of the biological assay used. (B) Results of 1-point screening at 10 μM of each compound using the cAMP assay. (C) Results of 3-point screening at the same concentration. Only compounds with an E_max_ > 50% in 1-point screening were selected. X-axis, numerical ID of compounds; Y-axis, results of cAMP assay (compared to DAMGO). All batches of experiments were validated with the Z' factor of DAMGO.

Several compounds demonstrated an efficacy greater than the theoretical E_max_ of the system, and these compounds were assumed to possess cell toxicity-related activities. Because the cAMP assay involves competition between intracellular cAMP and introduced cAMP, relative levels of introduced cAMP are increased when the intracellular cAMP level is suppressed due to opioid receptor activation. Hence, fluorescence signals will be stronger and signal quantification is greater. However, if compounds induce cell toxicity, intracellular cAMP levels can be depleted due to dysfunction of cell viability. Hence under toxicity conditions, the HTRF signal of the assay can be greater than that obtained with a full agonist. To identify whether the efficacy was due to receptor-ligand activation, the concentration-response curves of compounds were classified; the curves of toxic compounds are not sigmoidal (data not shown).

### Evaluation of concentration-dependent curves of compounds

We used a 4-parameter logistic curve model to analyze the concentration-dependent curve. The model consists of E_max_, E_min_ (basal response of cell system), EC_50_, and n (Hill slope). Ideal response curves should demonstrate a saturated maximal curve profile and a Hill slope between 0.7 and 1.5 (the Hill slope reflects the ligand-receptor binding mode). For Hill slopes markedly more different than 1.0, the ligand may bind at multiple sites on the receptor^32^. Moreover, similar E_max_ values and Hill slopes which approximate 1.0 are required to effectively compare pathways or compounds using relative activity^33^. Because Oliceridine and PZM21 are only partial agonists for OPRM^23^, we set the E_max_ criteria for discovering candidates as > 90% (compared to DAMGO) in order to discover full agonists for OPRM. The results for all 32 are presented in Table 1. Experimental data was fitted by Prism 8 software to estimate E_max_ and Hill slope parameters. An acceptable estimation of E_max_ and Hill slope was obtained for approximately 50% of the compounds. However, E_max_ and Hill slope values could not be calculated for 15 compounds (including **7)**. In most cases, these concentration-curves had either not reached the maximum asymptote or there was a weak dependency between concentration and efficacy (data not shown). Finally, eight compounds (**2, 3, 17, 20, 21, 22, 23, 32**) were selected (Compound **17** showed an efficacy around 80% and an appropriate Hill slope), and their concentration-curves are shown in Figure 4. The Hill slope for the selected compounds was ~ 1.0 (between 0.9 and 1.4), suggesting that these ligands bind at a single site on the target receptor.

**Figure 4.**
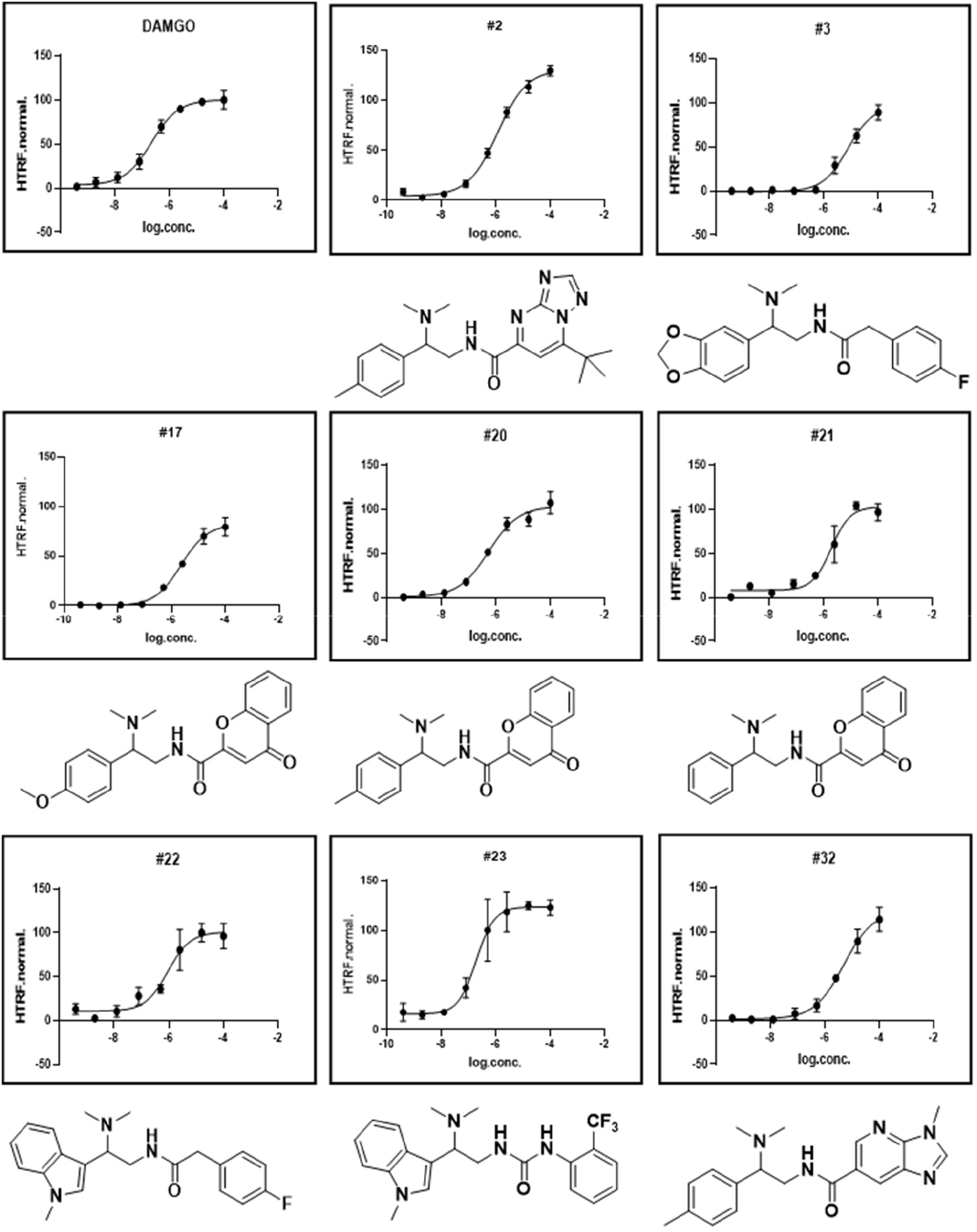
OPRM concentration-response curves of eight identified compounds and DAMGO. The data were normalized by the E_max_ of DAMGO. X-axis, log concentration of treated compounds; Y-axis, cAMP assay results normalized to DAMGO. All experiments were performed at least three times with triplicate-well replication (and confirmed using a Z' factor to obtain reproducible data). The structure of each compound is shown below each graph.

**Table 1.** E_max_ and Hill slope for 32 compounds with OPRM.

| ID | Curve E<sub>max</sub><br>(Normalization) | Hill slope | ID | Curve E<sub>max</sub><br>(Normalization) | Hill slope |
| --- | --- | --- | --- | --- | --- |
| 1 | 93.3 | 1.8 | 17 | 82.2 | 0.9 |
| 2 | 98.4 | 1.2 | 18 | 108.9 | 1.9 |
| 3 | 97 | 1.2 | 19 | 35.9 | 47.9 |
| 4 | 100 | 6.9 | 20 | 102 | 0.8 |
| 5 | 101.2 | 2.2 | 21 | 100.1 | 1.4 |
| 6 | 100 | uncalculated | 22 | 100.2 | 1.1 |
| 7 | uncalculated | uncalculated | 23 | 99.1 | 1.3 |
| 8 | 95 | 2.2 | 24 | uncalculated | -0.2 |
| 9 | uncalculated | uncalculated | 25 | uncalculated | uncalculated |
| 10 | uncalculated | 0.6 | 26 | uncalculated | uncalculated |
| 11 | uncalculated | uncalculated | 27 | uncalculated | uncalculated |
| 12 | 84.6 | 2.4 | 28 | uncalculated | uncalculated |
| 13 | 65.6 | uncalculated | 29 | uncalculated | 1.6 |
| 14 | 100 | uncalculated | 30 | 81.1 | 1.8 |
| 15 | 104 | 0.6 | 31 | uncalculated | uncalculated |
| 16 | uncalculated | 0.7 | 32 | 96.5 | 1.2 |

### Efficacy comparison of compounds of 2^nd^ library to G protein and β-arrestin2

Interestingly, all 8 compounds have the same scaffold, which is N-[2-(dimethylamino)propyl]formamide. The scaffold is shorter than PZM21 and their substituents are directly linked. Therefore, the new scaffold is expected to cover a different chemical space to PZM21, and to have a different pharmacological effect. To confirm the potential of this scaffold as a dual biased agonist, we conducted a substructure search in MolPort. In total, 53 second-round chemicals were purchased and tested. Following the flow chart shown in Figure 3A, we conducted experiments to evaluate the functional selectivity of all 53 compounds for the target receptors (OPRM and OPRD). Although a few biased agonists for OPRK have previously been reported^34,35^, OPRK was not selected as a target in the present study.

The efficacy of these compounds was evaluated using the cAMP assay (for G protein pathway) and the β-arrestin2 recruitment assay. E_max_ was estimated from eight-point (in triplicate) concentration-response curves. The results were normalized using full agonist (DAMGO for OPRM and SNC80 for OPRD). The E_max_ values are presented in Figure 5. Figure 5A and 5C report E_max_ values from the cAMP and β-arrestin2 assays, respectively. We first filtered compounds showing approximately 90% E_max_ (relative to reference agonist) for the G_i_ pathway and less than 15% E_max_ for the β-arrestin2 pathway. Twenty compounds including Oliceridine, PZM, and compounds ID110460001, ID110460002, and ID110460003 were selected. Using Z' factor values (≥ 0.5), we determined that 20 compounds had a reliable effect. We also confirmed that the concentration-response curves had a maximum asymptote with a Hill slope indicative of a sigmoidal curve (> 0.6). Compounds ID110460001, ID110460002, and ID110460003 were selected as final candidates. These compounds showed the functionality of a full agonist for the Gi pathway, and the functionality of a very weak partial agonist for the β-arrestin2 pathway (Figure 5D and Supplementary Figure S2). Furthermore, their concentration-response curves were sigmoidal (Supplementary Figure S3) like PZM21. Oliceridine was identified as a biased agonist for OPRM (but not OPRD), confirming the previous study^4^. Although Oliceridine behaved similarly to a full agonist in the G_i_ pathway, it showed a large distribution of variance and reduced low Hill slope (Table 2). Although experiments often yield Hill slopes slightly different to 1.0^32^, receptor-ligand reaction curves usually show a maximal asymptote. However, when the Hill slopes approach zero, maximal asymptotes disappear. Hence, Oliceridine may have multiple binding sites on OPRM, or it may simultaneously activate another pathway to affect the cAMP assay. In a previous study^4^, Oliceridine was evaluated by the relative method to calculate the effect of the log ratio of E_max_ and potency (EC_50_)^36^. In our study, these Oliceridine values were well-fitted to OPRM (Table 2 and Figure 5B). However, the method has limitations because it does not reflect the shape of the response curve (including the Hill slope). A new criteria to estimate efficacy is required. In this regard, the Hill slope may better report on the mode of action. As mentioned above, all three compounds identified are expected to show improved unity ligand binding-induced conformation changes in the receptor which increase intrinsic efficacy compared with Oliceridine.

**Figure 5.**
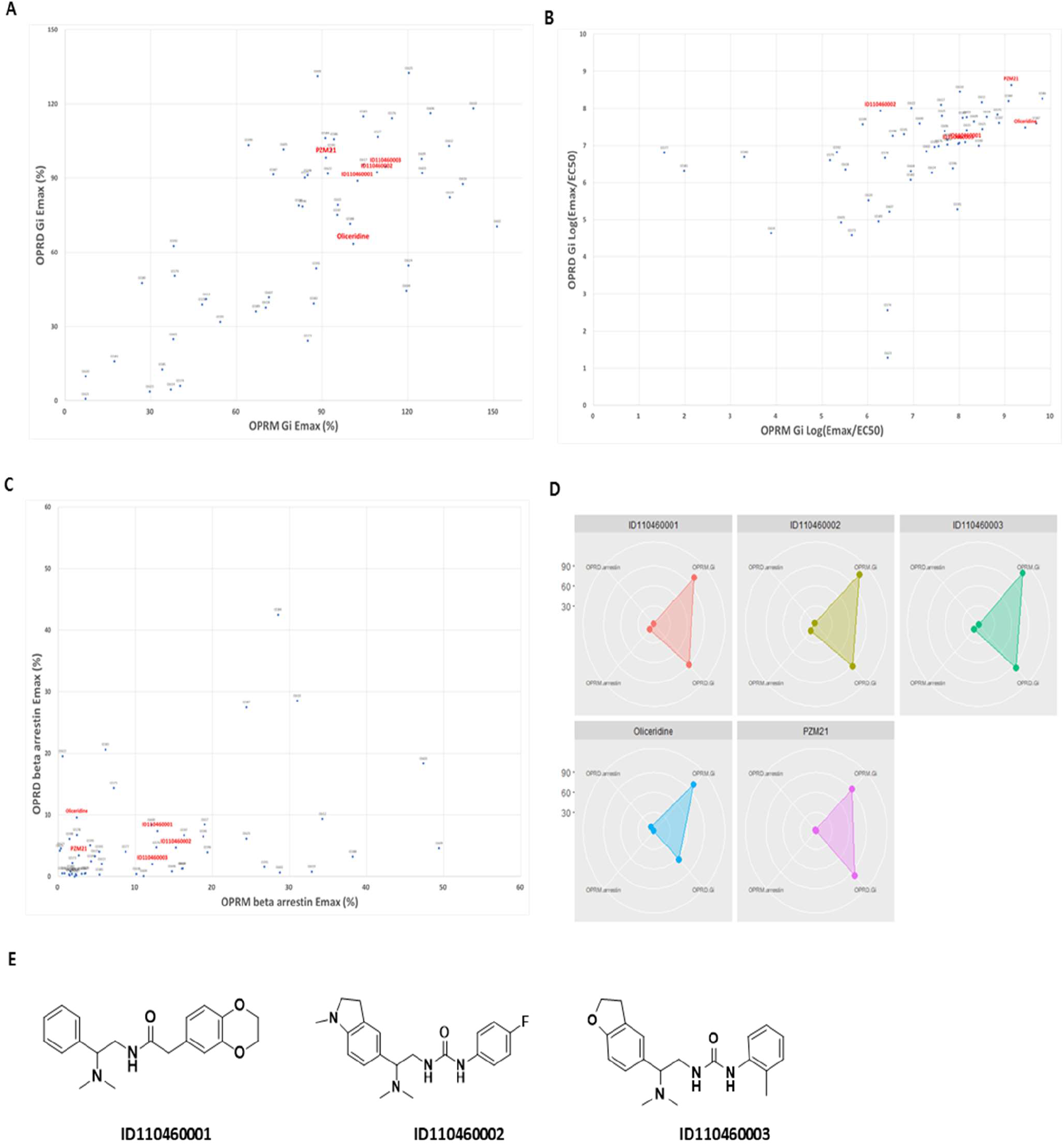
Evaluation of 2^nd^ library compound efficacy for the G protein and β-arrestin2 pathways. (A) cAMP assay. X-axis, mean efficacy on OPRM; Y-axis, mean efficacy on OPRD. (B) To compare relative efficacy, the log-ratio between E_max_ and EC_50_ was calculated. (C) β-arrestin2 recruitment assay. X-axis, mean efficacy on OPRM; Y-axis, mean efficacy on OPRD. (D) Radar chart of Oliceridine, PZM21, compound ID110460001, compound ID110460002, and compound ID110460003 using results from (A) and (C). (E) Structures of tested compounds. All experiments were performed at least three times (in triplicate). Data were normalized against full agonists (DAMGO for OPRM and SNC80 for OPRD).

**Table 2.** results of Oliceridine, PZM21, ID110460001, ID110460002, and ID110460003 on OPRM and OPRD.

|  | OPRM |  |  |  | OPRD |  |  |  |
| --- | --- | --- | --- | --- | --- | --- | --- | --- |
|  | Gi E<sub>max</sub> | Gi Hill slope | Gi Log<br>(Max/EC<sub>50</sub>) | β-arrestin 2<br>E<sub>max</sub> | Gi E<sub>max</sub> | Gi Hill slope | Gi Log<br>(Max/EC<sub>50</sub>) | β-arrestin 2<br>E<sub>max</sub> |
| Oliceridine | 100.8±30.5 | 0.4 | 9.1 | 2.4±1.1 | 63.4±16 | 1.3 | 8.6 | 9.6±16.2 |
| PZM21 | 91.3±10.1 | 0.7 | 9.5 | 2.7±0.8 | 98.3±6.8 | 0.8 | 7.5 | 3.4±3 |
| ID110460001 | 102.3±10.4 | 1.0 | 8.1 | 12.9±0.6 | 88.9±11.1 | 1.4 | 7.3 | 7.4±10.2 |
| ID110460002 | 109.1±10.2 | 0.9 | 7.8 | 15.3±3.6 | 92.4±18 | 1.4 | 7.9 | 4.7±0.8 |
| ID110460003 | 112.2±20.8 | 1.5 | 8 | 12.2±4.3 | 94.6±11.2 | 1.2 | 7.1 | 2±2 |

### Evaluation of intrinsic efficacy (versus excess receptor amplification)

As mentioned above, cell lines used to measure the efficacy of molecules are generally engineered to overexpress target receptors. Moreover, if experiments target downstream signaling, universal signaling factors (for instance cAMP, Ca^2+^, etc.) may be involved, increasing the probability that the results will be amplified. Indeed, several groups have suggested that bias calculations between cAMP and β-arrestin2 would have disproportionate results^13,23^. Thus, Oliceridine and PZM21 appear to be weak partial agonists in the G protein pathway in OPRM. To investigate whether the efficacy of our compounds (ID110460001, ID110460002, ID110460003) was affected by amplification, we conducted the Furchgott method^37^. When a fraction of the receptor population is permanently inactivated by an irreversible antagonist, the equiactive response of the agonist produces a specific pattern in their mechanism of action. If the agonist has high intrinsic efficacy, the E_max_ can resist decreases in receptor population. However, if the agonist showed maximal response while exploiting low intrinsic efficacy with the assistance of a rich receptor population, the E_max_ would be depressed^38^. β-Funaltrexamine hydrochloride (β-FNA) and β-Chlornaltrexamine dihydrochloride (β-CNA) were used as irreversible antagonists for OPRM and OPRD, respectively.

The OPRM results are presented in Figure 6, and the OPRD results are presented in Figure 7. As shown in Figure 6, DAMGO, a full agonist of OPRM, yielded an E_max_ of 64±1.8% at maximal concentration (10 nM) in FNA treatment. In contrast, Oliceridine and PZM21 showed maximal responses of 43±3.8% and 47±2.8%, respectively. This difference was significant (p<0.001) compared with DAMGO. Remarkably, ID110460001, ID110460002, ID110460003 maintained E_max_ values of 73.8±3.8%, 77.8±3.3%, and 70.7±2.9%, respectively (all higher compared with DAMGO). Thus, we confirmed that the efficacy of these three compounds was due to high intrinsic efficacy (and not amplification).

**Figure 6.**
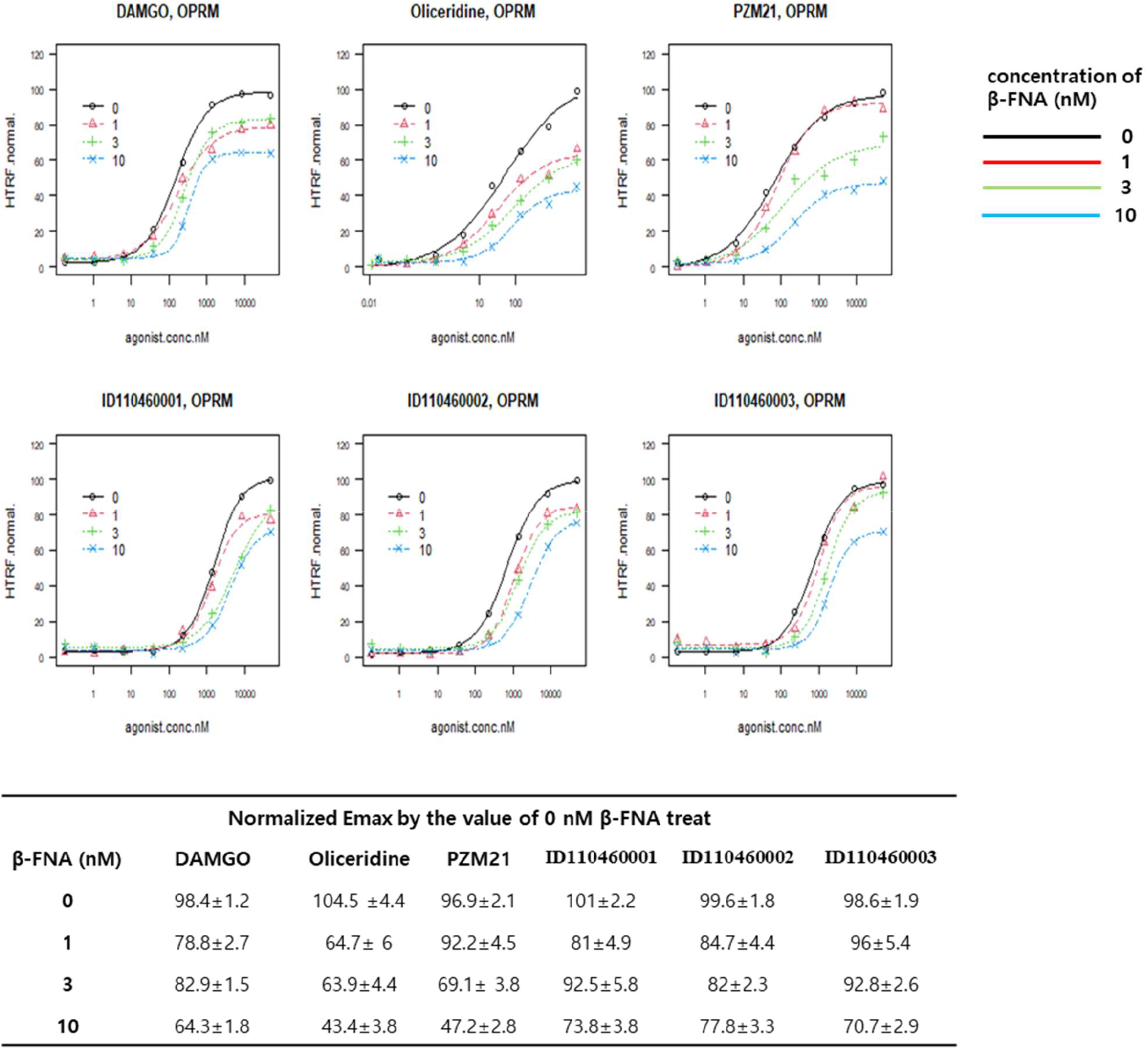
cAMP assay conducted in cell lines with a manipulated OPRM receptor population. DAMGO, Oliceridine, PZM21, ID110460001, ID110460002, and ID110460003 were applied to cell lines pretreated with various concentrations of β-FNA (an irreversible antagonist for OPRM). The changes in Emax due to β-FNA treatment are noted in the table below. Black, 0 nM β-FNA; Red, 1 nM β-FNA; Green, 3 nM β-FNA; Blue lines, 10 nM β-FNA. All experiments were performed at least 3 times (in triplicate).

**Figure 7.**
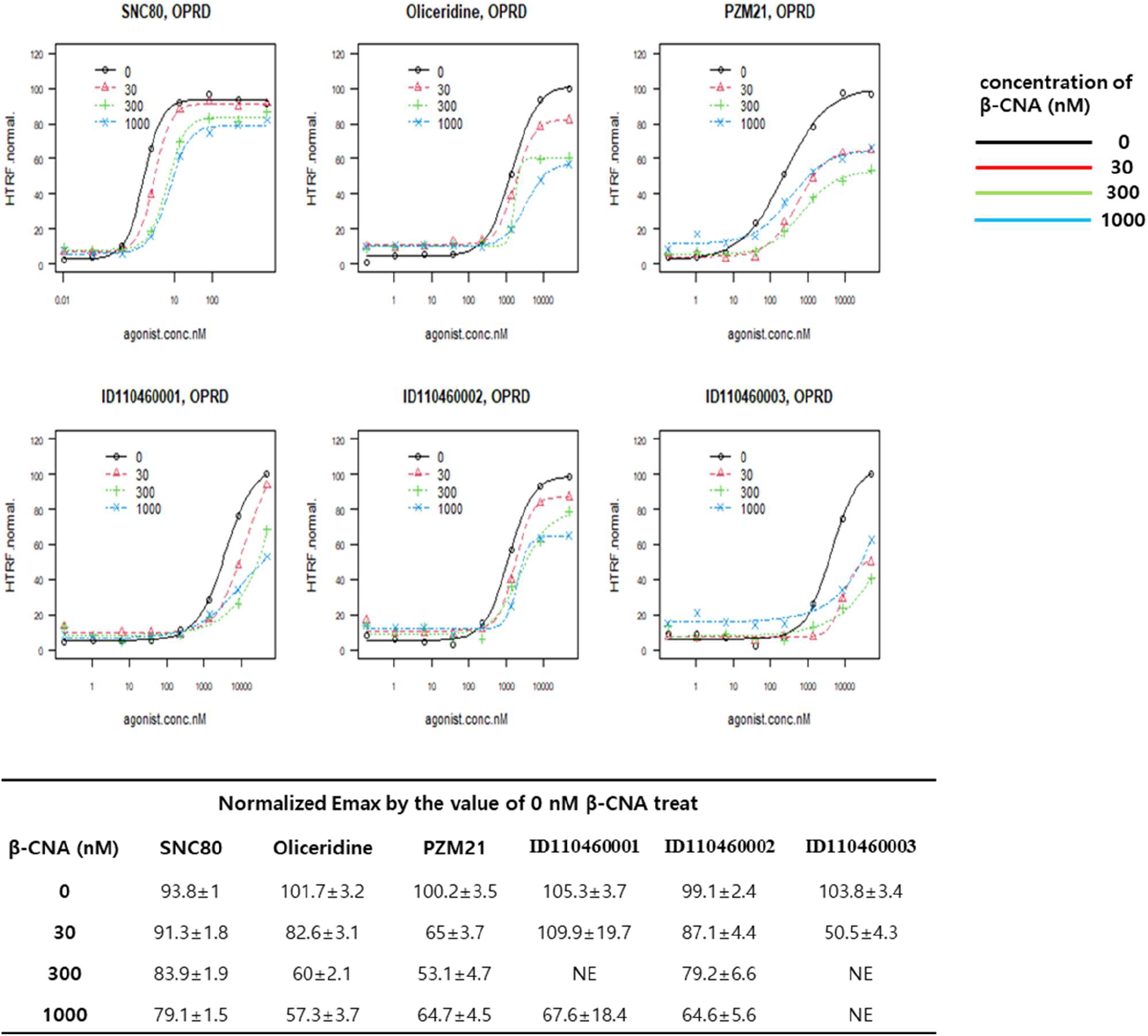
cAMP assay conducted in cell lines with a manipulated OPRD receptor population. SNC80, Oliceridine, PZM21, ID110460001, ID110460002, and ID110460003 were all applied to cell lines pretreated with various concentrations of β-CNA (an irreversible antagonist for OPRD). The changes in Emax due to β-CNA treatment are noted in the table below. Black, 0 nM β-CNA; Red, 30 nM β-CNA; Green, 300 nM β-CNA; Blue lines, 1000 nM β-CNA. All experiments were performed at least 3 times (in triplicate).

As shown in Figure 7, SNC80, a full agonist of OPRD, maintains an E_max_ up to 79.1±1.5%. The maximal response of Oliceridine and PZM21 was decreased to 57.3±3.7% and 64.7±4.5%, respectively. Hence, these two compounds act as partial agonists to OPRD (not full agonists). In contrast to the results observed with OPRM, ID110460001, ID110460002 and ID110460003 failed to maintain the maximum response to OPRD (compared with SNC80). The response curves of ID110460001 and ID110460003 lost the maximum asymptote, indicating that they were affected by the number of receptors. Consequently, the potency of drugs was attenuated. Although compound ID110460002 maintained the maximal asymptote of the response curve, the maximal response was similar to PZM21 (64.6±5.5%) and significantly less than SNC80. Compound ID110460002 may behave as a partial agonist similar to PZM21 when binding OPRD.

From the above results, we can confirm our compounds shows an OPRM-specific mechanism of bias. Because compounds demonstrate high intrinsic efficacy (and was not affected by the receptor population), we expect that compounds will act as a biased agonist for the G protein pathway of OPRM *in vivo*. However, the efficacy of compounds for OPRD is affected by receptor population, and ID110460002 possesses better profile than others. Therefore, we expect that ID110460002 would show different properties depending on the tissue or organs (due differences in receptor expression levels). OPRM is a major target receptor of morphine, and the goal of biased agonists is pain relief without side effects. Therefore, the properties of compound ID110460002 are of particular interest. In future studies, we plan to test the efficacy of ID110460002 in animal models.

### Analysis of binding modes

To obtain a detailed insight into receptor-ligand binding, a human OPRM homology model was built using the published mouse OPRM crystal structure (5C1M.pdb). We subsequently performed a docking study using this human OPRM homology model and the human OPRD X-ray crystal structure (PDB id: 6PT2), analyzing the interactions between receptor and hit compounds. The predicted binding modes of compounds ID110460001, ID110460002 and ID110460003 with OPRM are shown in Figure 8.

**Figure 8.**
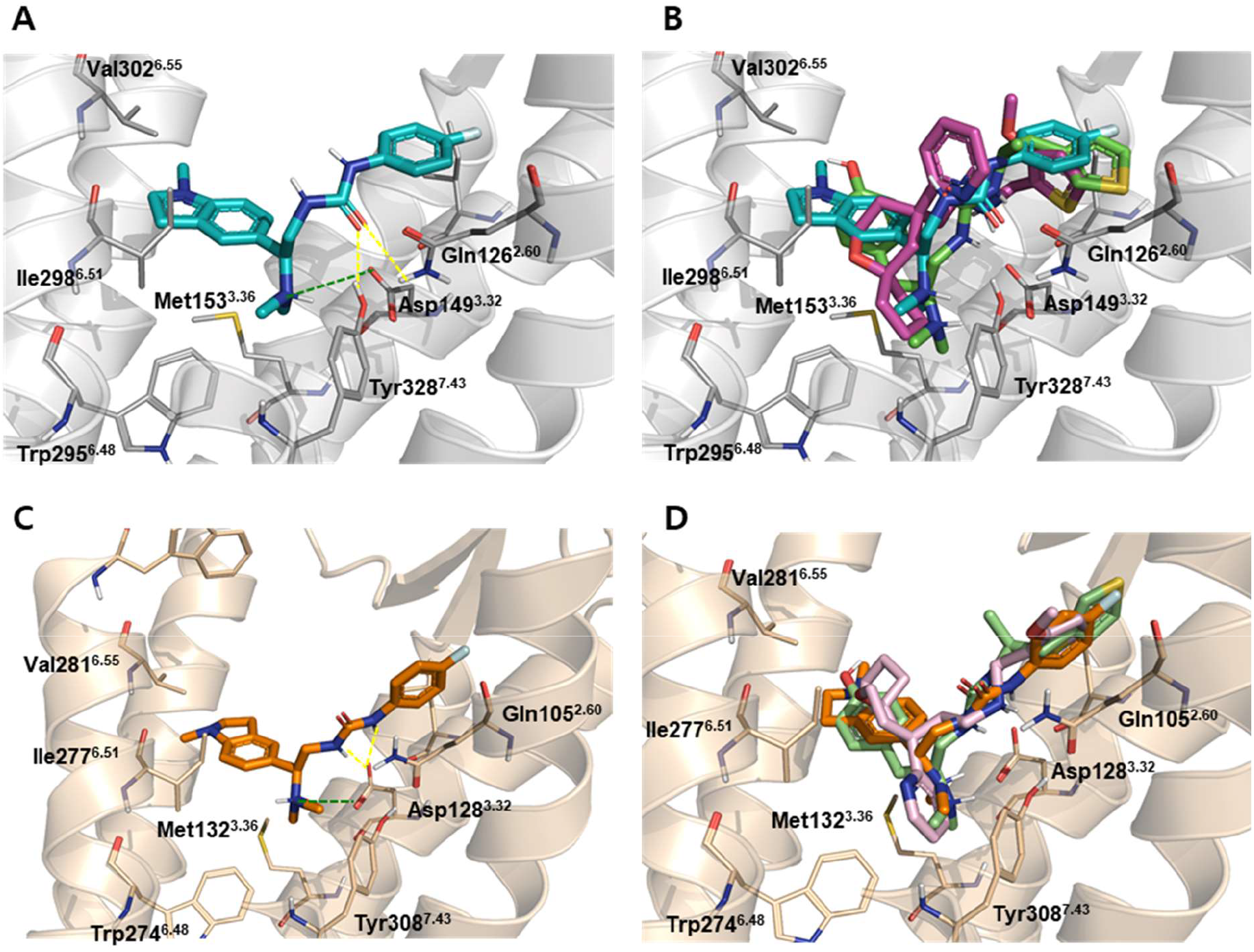
Predicted binding modes of biased agonists for OPRM and OPRD. Binding modes of (A) ID110460002 (teal), and (B) PZM21 (green), Oliceridine (magenta) in active OPRM. Binding modes of (C) ID110460002 (orange), and (D) PZM21 (light-green), Oliceridine (light-pink) in active OPRD. Hydrogen bonds are shown as yellow dashed lines, and the important salt bridge is depicted by the green dashed line.

The protonated nitrogen atom of these compounds formed a strong salt bridge with Asp149^3.32^ (superscripts indicate Ballesteros-Weinstein numbering), which is a well-known anchoring point for binding of all opioid receptors and modulators. In all interactions between OPRM and ID110460001, ID110460002 and ID110460003 the N-dimethyl group was involved in π-cation interactions with Tyr328, while the urea carbonyl formed hydrogen bonds with Gln126 and Tyr328. Furthermore, hydrophobic interactions with Ile298 and Val302 were also conserved. In addition to these common interactions, the methylindoline group of ID110460002 was involved in specific hydrophobic interactions with Val238 and His299, and the fluorobenzene was involved in specific hydrophobic interactions with Ile146 and Asp56 (Figure 8A). The benzene ring of ID110460001 and the 2,3-dihydro benzofuran of ID110460003 also formed π-sulfur interactions with Met153 (Supplementary Figure S1).

In the docked configuration, ID110460002, PZM21, and Oliceridine, all maintained the key ionic interaction with Asp149, and all three contacted the orthosteric binding pocket (Figure 8C). However, in contrast with our three hit compounds, PZM21 and Oliceridine did not make hydrogen bonds with Gln126 and Tyr328. Moreover, PZM21 and Oliceridine also failed to make contact with Val302, which is involved in well-known hydrophobic interactions. Therefore, the binding modes of the hit compounds differed significantly from those of PZM21 and Oliceridine, and this is reflected in the pharmacological effects shown in Figure 6.

As shown in Figure 8C and 8D, the docked pose of ID110460002 bound to OPRD (6PT2.pdb) is similar to that of PZM21 and Oliceridine. In all three cases, the key ionic interactions between cationic amine and Asp128 were present, as were hydrophobic interactions involving Tyr129 and Val217. Furthermore, the urea group of ID110460002 and PZM21 were involved in hydrogen bonding interactions with Asp128. As shown in Supplementary Figure S1, ID110460001, ID110460002, ID110460003, PZM21, and Oliceridine all showed similar binding modes for OPRD.

### Conclusions

Because OPRM and OPRD play an important role in pain, these receptors are promising molecular targets. Here, we report the discovery of compound ID110460002, an OPRM, OPRD dual biased agonist. Remarkably, ID110460001, ID110460002 and ID110460003 are all improved biased agonists for OPRM (compared with Oliceridine and PZM21). Although Oliceridine and PZM21 both show biased efficacy toward the G protein pathway (compared with the β-arrestin2 pathway), this is a result of cAMP signal amplification in the G protein pathway. Hence, Oliceridine and PZM21 are not suitable as clinical biased agonists. However, biased efficacy to the G protein pathway of OPRM has been confirmed to be intrinsic for ID110460001, ID110460002 and ID110460003. Therefore, these three compounds may represent leads for a physiological real biased agonist to manage pain without side effects. On the OPRD pathway, maximal efficacy of ID110460002 toward the G protein pathway is higher than for Oliceridine. However, ID110460002 is affected by receptor population. While an improved mechanism of action specifically for OPRD was not demonstrated, ID110460002 would be expected to show better *in vivo* efficacy because it has a higher E_max_ (compared with Oliceridine).

In this study, virtual screening and docking studies were performed to construct a compound library and subsequently investigate binding modes between receptors and compounds. Using virtual screening, we efficiently constructed a compound library and identified a number of candidates. Through the docking study, we could investigate the different binding modes of these compounds for OPRM and OPRD. As a result, we have high expectations that we can now develop the first biased agonist of opioids from compound ID110460002.

## Materials & Method

### Materials

Oliceridine was purchased from AdooQ (CA, US). PZM21 was purchased from MedKOO bioscience (NC, US). SNC80 and [D-Ala (2), NMe-Phe^4^, Gly-ol (5)]-enkephalin (DAMGO) were purchased from Tocris (Bristol, UK). Test compounds were purchased from MolPort (Riga, Latvia). β-Funaltrexamine hydrochloride (β-FNA) and β-Chlornaltrexamine dihydrochloride (β-CNA) were purchased from Merck (Darmstadt, Germany). The PathHunter β-arrestin assay kit was purchased from DiscoverX (CA, USA). The cAMP assay kit was purchased from Cisbio (codolet, France). Lipofectamine 2000 was purchased from (Thermo Fisher Scientific (MA, US).

### Construction of μ-opioid or δ-opioid receptor overexpressed cell lines

Human Embryonic Kidney (HEK)-293 PathHunter EA Parental Cell Lines stably expressing a β-arrestin2 - β-galactosidase fragment (EA) fusion were purchased from DiscoverX (CA, US). The human OPRM1 gene (NM00914.3) and human OPRD1 gene (NM00911.3) were cloned into a pCMV-ProLink plasmid containing a complementary β-galactosidase fragment purchased from DiscoverX ^4^. HEK-293 cells were transfected with pCMV-ProLink plasmids expressing a receptor – complementary β-galactosidase fragment fusion to generate stable cell lines. Cells were grown in Dulbecco's Modified Eagle's Medium (DMEM) containing 10% fetal bovine serum, 500 μg/mL G418, and 200 μg/mL Hygromycin B.

### Measurement of intracellular cyclic adenosine monophosphate (cAMP)

cAMP measurements were conducted to investigate G protein pathway signal activation. cAMP-homogenous time-resolved fluorescence kits were used for analysis. OPRM/ OPRD bind Gαi protein upon receptor activation, inhibiting intracellular cAMP accumulation induced by forskolin^4^. Therefore, low cAMP levels are indicative of opioid receptor activation. To prevent degradation of intracellular cAMP, all experiments were conducted in the presence of 500 μM 3-isobutyl-1-methylxanthine (IBMX). The test compounds were diluted with dimethylsulfoxide (DMSO), at a maximum final concentration of 1% (to prevent cell membrane disruption), or phosphate-buffered saline (PBS)^25^. Irreversible antagonists (β-FNA for OPRM, β-CNA for OPRD) were used to reduce the population of receptors on the cell surface to diminish signaling amplification^26^. Cells were pre-treated with various concentrations of irreversible antagonists for 20 minutes before cAMP measurement. All data were measured using a FlexStation 3 (Molecular Devices, CA, US) and normalized to the maximal assay response obtained using a full agonist (DAMGO for OPRM, SNC80 for OPRD). In assays using irreversible antagonist pre-treatment, data were normalized to the maximal response of test compounds without pre-treated irreversible antagonist group.

### β-arrestin2 recruitment assay

After receptor activation, the β-arrestin2 fused with β-galactosidase fragment binds to the c-terminus of the receptor-complementary fused with β-galactosidase fragment. Thus, a functional β-galactosidase is reconstituted. The kinase reaction was detected by chemiluminescence using the PathHunter assay. Identical cell lines were used in both the β-arrestin2 assay and the cAMP assay. Again, data were measured using the FlexStation 3 and normalized as above.

### Ligands and database preparation

The KCB chemical library (comprising 430,000 entries) was preprocessed to eliminate reactive compounds and inorganic admixtures, and assess Lipinski’s rule of five and PAINS patterns using the Pipeline Pilot ^27^. The low-energy 3D structures of compounds were generated by LigPrep using the default settings ^28^. In total, 53 compounds were purchased from Mol-Port (http://www.molport.com) for testing.

### Pharmacophore model

The conformers of three biased agonists were generated by Discovery studio using the default parameters. Using the lowest conformers, we constructed a pharmacophore model by Phase implemented in Schrodinger ^28^. The pharmacophore model was developed using a set of pharmacophore features to generate sites for all three compounds. The alignment was measured using a survival score with default values for the hypothesis generation.

### Docking

Mouse OPRM (PDB id: 5C1M) ^29^, active state human OPRD (PDB id: 6PT2) ^30^, and human OPRM homology models were prepared using the Protein Preparation Wizard Workflow provided in the Maestro module of Schrodinger Suite 2019-4. A receptor grid was generated covering a 20 Å region centered at the co-ligand of the complex structure. Low-energy 3D structures of ligands were docked using Glide in the Standard Precision (SP) mode (using default values). Protein-ligand interactions were analyzed using Discovery Studio Modeling Environment v4.026 and the docking models were displayed using PyMOL version 2.0.47.

### Homology model

The mouse OPRM X-ray structure (PDB id: 5C1M) was selected as a template structure for human OPRM. The amino acid sequence of human OPRM (accession number: P35372) was retrieved from the UniProtKB database (http://www.uniprot.org). The target and template sequences (sequence identity, 93.75%) were aligned using the ClustalW method in Prime, followed by manual adjustment for gaps in the transmembrane region. A 3D homology model of human OPRM was built using Prime in Schrödinger Suite 2019. The overall stability of the protein was assessed using the Ramachandran plot. Other procedures were as described in the docking study for human OPRM.

### Statistical analyses

Data analysis and statistical hypothesis tests were conducted using Prism 7 version (GraphPad Inc., CA, US) or R software 4.0.3 version (R Core Team, Vienna, Austria). We used Analysis of variance with a Tukey-Kramer multiple comparison *posthoc*-test or means of Z-test as reported by Ritz et al^31^. The E_max_ (maximal response of cell system), E_min_ (minimum response of cell system), hill slope, and EC_50_ (drug concentration at half-maximal response) of compounds were fitted using a 4-parameter logistic curve equation. The efficacy data of all small molecules were qualified by the Z’ factor. The efficacy data sets were considered suitable if the Z’ factor exceeded 0.5 (and discarded otherwise).

## ASSOCIATED CONTENT

Supplementary Figure S1; Supplementary Figure S2; Supplementary Figure S3.

## AUTHOR INFORMATION

## Author Contributions

*Conceptualization*: D.S.L.. *Visualization*: J.H.L; D.S.L.. *Methodology*: J.H.L; S-Y.S.. *Investigation*: J.H.L; S-Y.S; W.J.; S-J.H; J.B.. *Formal analysis*: J.H.L.. *Validation*: S-Y.S.. *Writing*: J.H.L; D.S.L. *Supervision*: D.S.L.. *Project administration*: D.S.L.. All authors have given approval to the final version of the manuscript.

## ACKNOWLEDGMENT

The chemical library used in this study was kindly provided by Korea Chemical Bank (www.chembank.org) of the Korea Research Institute of Chemical Technology.

