## Supplemental Figures for "Discovery of μ, δ-Opioid receptor dual biased agonists that overcome the limitation of prior biased agonists"

### Supplementary Information

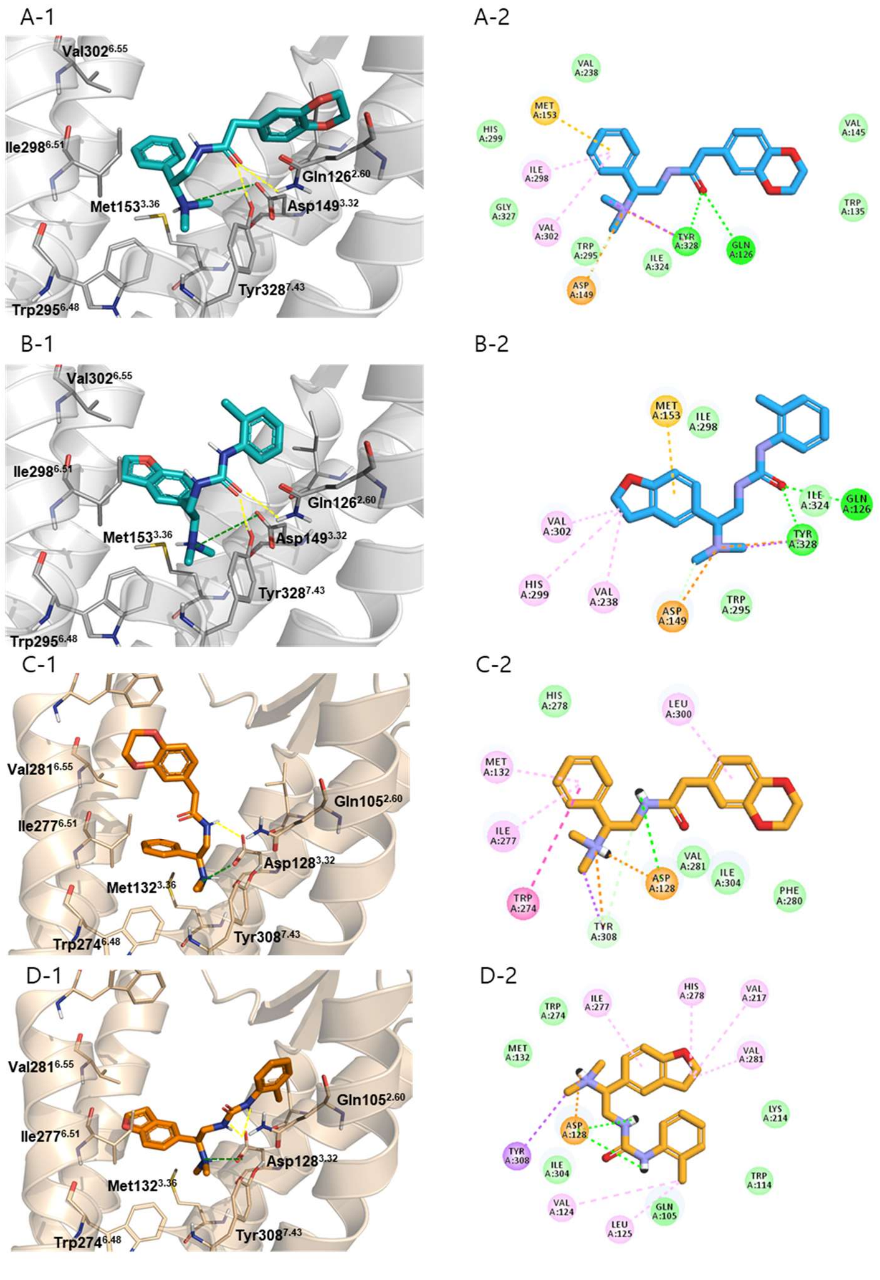


**Figure S1. Predicted Binding modes of biased agonists for OPRM and OPRD.** Binding modes of (A) ID110460001 (teal) and (B) ID110460003 (teal) in active OPRM. Binding modes of (C) ID110460001 (orange) and (D) ID110460003 (grange) in active OPRD. Hydrogen bonds are shown as yellow dashed lines and the salt bridge is depicted as a green dashed line.

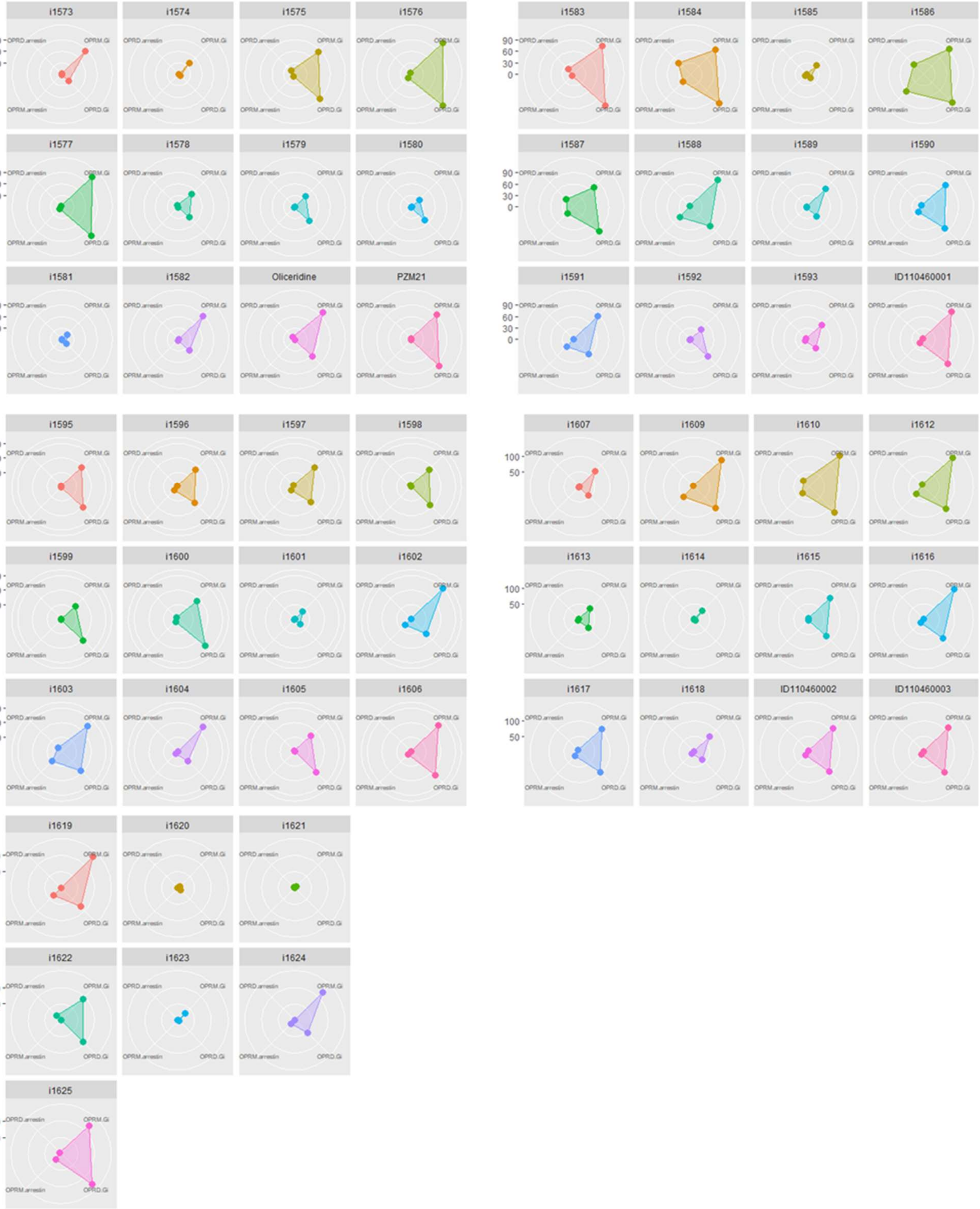


**Figure S2. Radar chart of 53 compounds.** The Radar chart presents the results of the cAMP assay and beta-arrestin 2 recruitment assay. The four vertices represent individual assay results for OPRM and OPRD; an idealized square graph represents OPRM, OPRD dual full agonists.

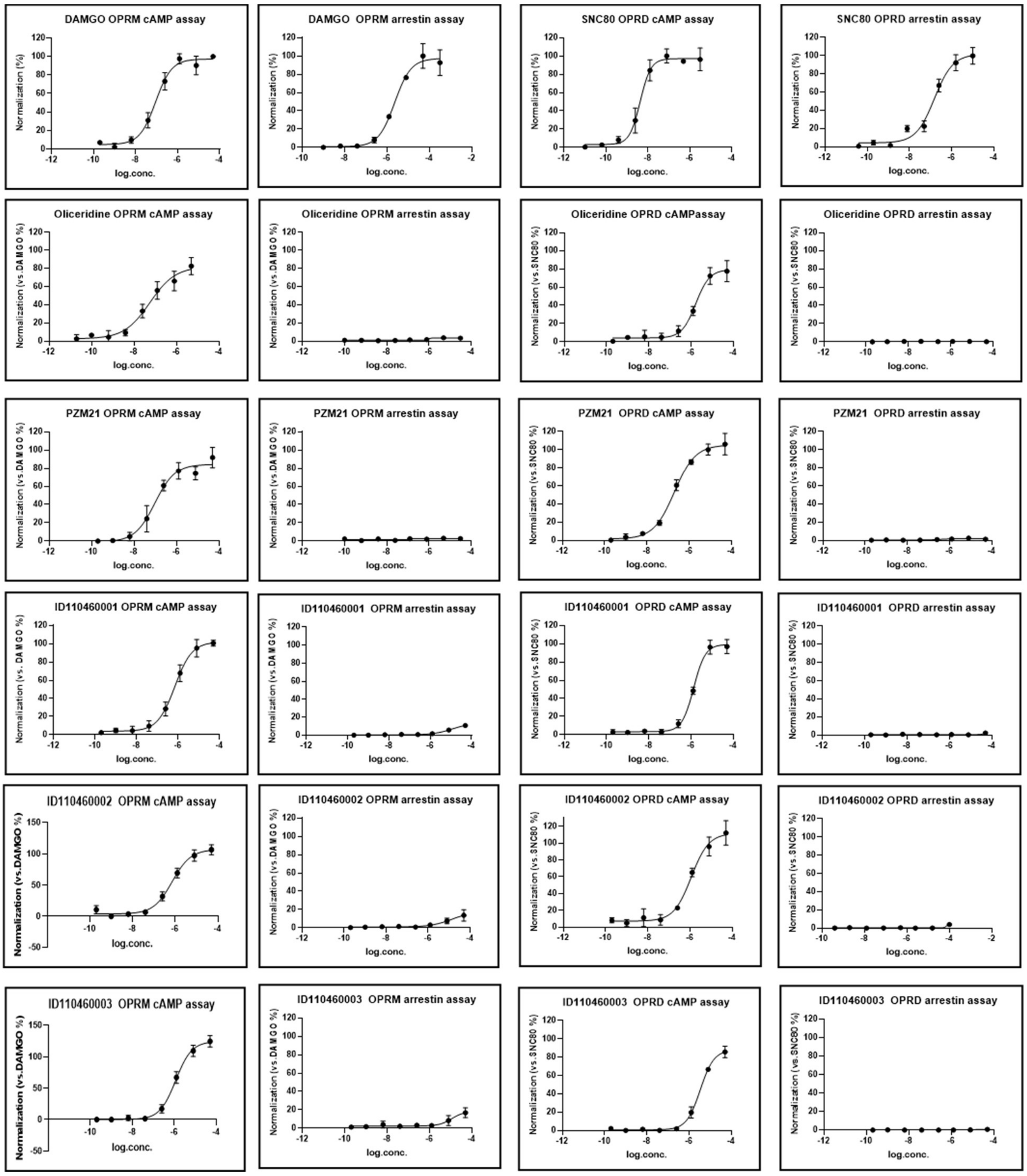


**Figure S3. OPRM and OPRD concentration-response curves of cAMP assay and beta-arrestin 2 recruitment assay results for five compounds (Oliceridine, PZM21, ID110460001, ID110460002, and ID110460003) and full agonist (DAMGO or SNC80)**. The OPRM and OPRD data were normalized by $E_{max}$ of DAMGO and $E_{max}$ of SNC80, respectively. X-axis, log concentration of treated compounds; Y-axis, assay results normalized by full agonist. All experiments were conducted (at least) three times with triplicate well replication, and results were confirmed using the Z' factor to obtain reproducible data.
